# Larvicidal activity of *Azadirachta indica, Melaleuca alternifolia*, and *Carapa guianensis* oil compounds and Carica papaya fermented extract on *Aedes aegypti* Larvicidal activity of oil compounds on *Aedes aegypti*

**DOI:** 10.1101/2020.05.18.102442

**Authors:** Sandra M. Torres, Luis A.R. Lima, Maria do Carmo A. Lima, Lêucio C. Alves, Valdemiro A.S. Júnior

## Abstract

The aim of this study was to evaluate the structural changes of *Aedes aegypti* larvae submitted to treatment with a product based on *Azadirachta indica, Melaleuca alternifolia, Carapa guianensis* oils and *Carica papaya* fermented extract. The larvae were exposed for 24h to the experimental groups: product formulated in concentrations of 50% (G1), 25% (G2), and 12.5% (G3); and negative control groups based on water (CN1) and dimethyl sulfoxide (CN2). By the end of the experimental period, some larvae were fixed in 4% buffered glutaraldehyde solution to be processed for optical microscopy. Larvae exposed to G2 and G3 presented more structural damage of the mesentery, Malpighi tubules and nerve ganglia. We conclude that the product formulated in 12.5% and 25% concentrations can be used in the population control of the 3rd larvae stage of *Aedes aegypti* by causing lethal injuries and avoiding the larvae development.

## Introduction

The dengue virus (DENV), an arbovirus transmitted by mosquitoes, became a major threat to human life in the Americas, reaching up to 23 million cases from 1980 to 2017. Brazil is among the most affected countries, presenting 13.6 million cases over the years (Salles et al., 2018). The *Aedes (Stegomyia) aegypti* is a diptera of great epidemiologic importance due to its role as dengue, yellow fever, Zika and Chikungunya virus transmitter (LEE et al., 2018; ZOHDY et al., 2018; PROW et al., 2018; ALVAREZ COSTA et al., 2018). Decreasing adults’ population density by using larvicides represents one of the most common ways of controlling this transmitter vector, since larvae are more accessible due to its low mobility and pre-established habitat (FILLINGER et al., 2011).

Larvicides obtained from botanic compounds, such as vegetable oils and essential oils and oleoresins, are an excellent alternative to the synthetic insecticides due to the variety of bioactive compounds obtained from renewable sources and rapid biodegradability, without the accumulation of toxic wastes in food or environment (ROEL, 2001; NAVARRO-SILVA et al., 2009; PAVELA, 2015).

Studies on the toxicity of several plant extract on *A. aegypti* larvae have been focusing on structural evaluations to understand their action sites (GUSMÃO et al., 2002; Arruda et al., 2003; 2003a). These studies usually aim to improve the knowledge regarding these products so it can be used later in the development of commercial insecticides (GUSMÃO et al., 2002; VALOTTO et al., 2010; BORGES et al., 2012). Thus, here we aim to evaluate the structural changes of larvae of *Aedes (*Stegomyia) *aegypti* (Diptera: Culicidae) submitted to treatment with a product formulated from *Azadirachta indica, Melaleuca alternifolia, Carapa guianensis* oils and *Carica papaya* fermented extract.

## Material and Methods

### Obtaining the formulated product

The formulated product was provided by Gued’s Biotecnologia^®^, and was composed by 1% of fixed seed oil from *Azadirachta indica*, 0.3% of fixed oil from *Melaleuca alternifolia’s* fruit, 1% of essential oil from *Carapa guianensis*, and 5% of bacterial fermented extract from *Carica papaya’s* fruit.

### Larvicidal assay

The toxicological test using third stage larvae was adapted from the methodology preconized by the World Health Organization (1970), with the modifications described as following (Jang et al., 2002). The test was carried out in triplicate with 900 larvae for each experimental group, totalizing 6300 specimens. The larvae were exposed to the solutions for 24 hours, being monitored every hour. The experimental groups followed the distribution found in Table 1. G1, G2 and G3 obtained mortality rates of 65%, 50% and 78%, respectively (Torres et al., 2014).

**Table 1:** Distribution of experimental groups of *Aedes aegypti* larvae exposed to the formulated product for 24 hours.

| Treated groups | Positive control | Negative control |
| --- | --- | --- |
| G1 – formulated product 50% (n=900) | CP1 – BTI CL<sub>90</sub> 0.37ppm<br>(n=900) | CN1 - dechlorinated water<br>(n=900) |
| G2 – formulated product 25% (n=900) | CP2 – BTI CL<sub>50</sub> 0.06ppm<br>(n=900) | CN2 - dimethyl sulfoxide<br>(n=900) |
| G3 – formulated product 12,5% (n=900) |  |  |

### Structural analysis

For the larvae histological study, (dead or live) specimens from all experimental groups were selected, fixed in 4% glutaraldehyde solution with 0.01M sodium phosphate buffer (PBS) of pH 7.3 and subsequently submitted to inclusion in historesin, according to Arruda et al. (2003). The 4 µm thick slices of the midgut, Malpighi tubule and nerve ganglia of *Aedes aegypti larvae* were stained in 1% toluidine blue and analyzed in optical microscope DM500E Leica© (Germany) in 400X magnification.

### Height and diameter of Malpighi tubule

The 3rd stage larvae were cross sectioned to measure the height of the epithelium and the diameter of the Malpighi tubules (µm), which were obtained through image capture using a Leica © DM500E optical microscope (Germany) in 400X magnification connected to a microcomputer. The methodology was adapted from Lastre et al. (2016), and the measurements were obtained from 15 tubules chosen at random. The epithelial height was measured using the diametrically opposite measurements of the basal membrane and the tubular lumen as reference. The diameter was measured considering the opposite basal membrane. All photomicrographs were evaluated through the software ImageJ version 1.52v (Image Analysis Software), using a linear reticular micrometer with 400X magnification.

### Statistical analysis

The data obtained were expressed through centrality and dispersion measurements (average ± standard deviation), following non-Gaussian distribution using the Mann-Whitney test or the Kruskal-Wallis non-parametric test, with Dunn’s post-hoc. The analyses were performed in the software InStat (GraphPad Software, Inc., 2000) The statistical treatment was designed with a significance level of p <0.05.

## Results

The structural changes found in the larvae exposed to the formulated product based on *Azadirachta indica, Melaleuca alternifolia*, and *Carapa guianensis* oils and *Carica papaya* fermented bacterial extract are shown in Figure 1.

**Figure 1:**
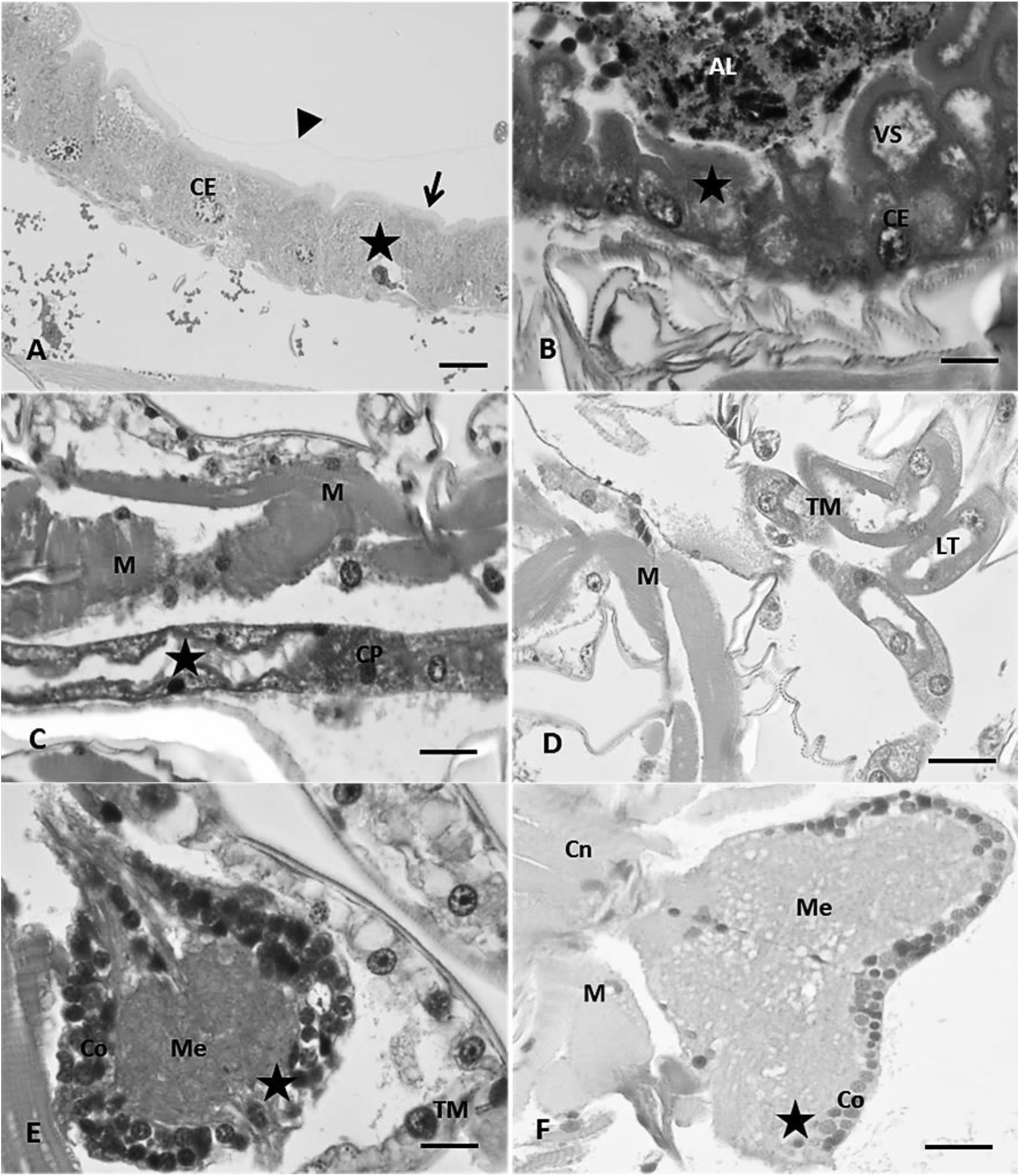
Photomicrography of intestine, Malpighi tubule and ganglia of third stage larvae of *Aedes aegypti* treated with formulated product based on *Azadirachta indica, Melaleuca alternifolia*, and *Carapa guianensis* oils and *Carica papaya* fermented extract, in 50%, 25% and 12.5% concentration. A: Midgut region of G1 with necrosed epithelial cells. 4µm Bar. B: Posterior region of the intestine of G2 with necrosis and cytoplasmatic vacuolization. 50µm Bar. C: Malpighi Tubule (TM) of G2 with lumen increase and tubular and muscular (M) necrosis. 50µm Bar. D: Malpighi Tubule of G3 with lumen increase and tubular necrosis. 50µm Bar. E and F: Ganglia of G2 and G3 with cortical necrosis and consequent cellular rarefaction and medullar vacuolization. 50µm Bar. Epithelial cell (CE); Secretion vesicle (VS); Food (AL); Cellular necrosis (star); Muscle (M); peritrophic matrix (arrow head); brush border (arrow); Main cell (CP); tubular lumen (LT); cortical region (Co); medullar region (M); nerve cord (Cn).

### Changes in Mesentery

In the mesentery of the G1 was observed a degenerative process of the epithelium, with cell swelling and cytoplasmic vacuoles formation, besides intestinal cells starting a process of necrosis. There was no peritrophic matrix and brush border. In G2, the intestinal injuries were the same, but a decrease in brush border height and maintenance of the peritrophic matrix was also observed, besides multifocal necrosis. The intestinal injuries were bigger in G3 when compared to G1 and G2. There was necrosis of extensive areas of the intestine, increased sub peritrophic space, extrusion of the peritrophic matrix, decrease of the brush border with signs of destruction and vacuolized secretory vesicles, and invagination of epithelial cells leading to an increase in the intercellular space.

The structural changes of the mesentery of *A. aegypti* larvae exposed to the industrial larvicide *Bacillus thuringiensis israelensis* (BTI) in CL_90_ 0.37ppm and CL_50_ 0.06ppm are shown in Figure 2.

**Figure 2:**
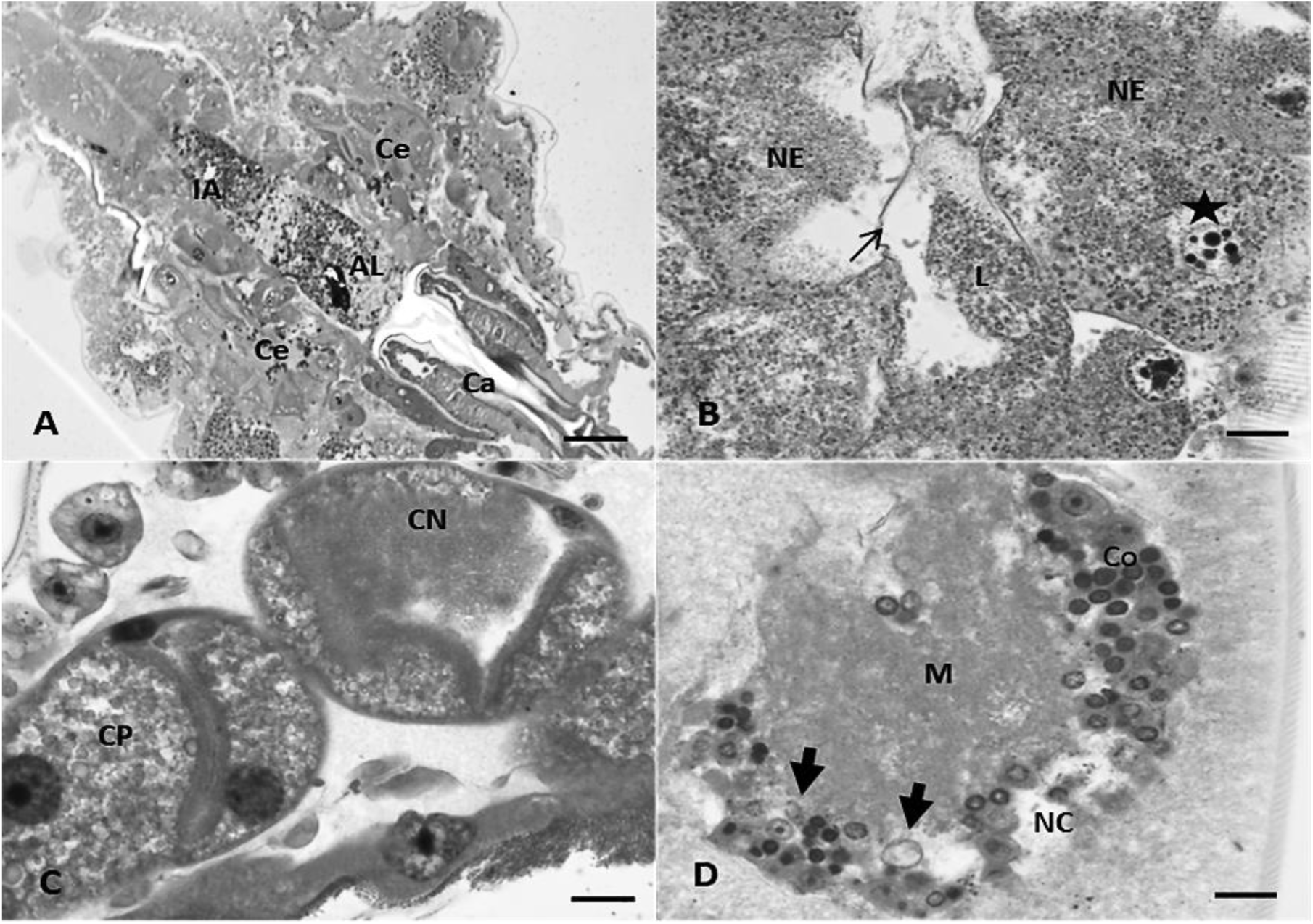
Photomicrography of intestine, Malpighi tubule and ganglia of third stage larvae of *Aedes aegypti* from the positive control group, treated with BTI CL_90_ 0.37ppm and CL_50_ 0.06ppm. A: Anterior region of the intestine presenting the extension of the necrosis. 200µm Bar. B: Closer observation of the necrosed midgut. 4µm Bar. C: Necrosed Malpighi tubule. 4µm Bar. D: Malpighi Tubule with cortical necrosis. 4µm Bar. Anterior intestine (IA); Gastric caecum (Ce); Cardia (Ca); Food (AL); Epithelial necrosis (NE); Intestinal lumen (L); Peritrophic matrix (thin arrow); Cortical necrosis (Thick arrow); Apoptotic bodies (star); Main cell (CP); Cellular necrosis (CN); Cortical region (Co); Medullar region (M).

The injuries in the epithelial cells of the intestine were observed throughout the organ, but the most relevant changes were found in the anterior mesentery, near the caecum and in the midgut. Intestinal wall and basement membrane diffuse necrosis and cell boundary loss with leakage of intestinal contents to the hemolymph were the most observed changes. The lesions were smaller in the posterior intestine, but still with cell necrosis, cytoplasmatic vacuolization, loss of secretory vesicles, peritrophic matrix, and brush border. Changes in the architecture of the mesenteric cells were not found in the negative control group.

### Malpighi tubule

Structurally, an increase in the Malpighi tubule’s lumen, necrotic cells, and cell and nuclear swelling with brush border maintenance can be seen in G1 (Figure 1). In G2, there was an extensive area of tubule necrosis, with increased tubular lumen and decreased brush border. Finally, G3 presented the same changes of G2, with an increase in vacuoles of cytoplasmic secretion and tubular lumen.

The diameter of the Malpighi tubules and the height of the epithelium of all experimental groups are described in Table 2. G1, G2 and G3 presented a dose-dependent decrease of the height (p<0.0001) in comparison with positive and negative control groups. However, the groups treated with the compound obtained the smallest tubular diameter, with G3 being the smallest among them (p = 0.0001) in comparison with G1 and G2.

**Table 2:** Height and diameter of Malpighi tubules (Average ± Standard Deviation) of *Aedes aegypti* larvae treated with product formulated from *Azadirachta indica, Melaleuca alternifolia*, and *Carapa guianensis* oils and *Carica papaya* bacterial fermented extract for 24 hours.

| Parameters ($\mu\text{m}$) | G1 | G2 | G3 | CP1 |
| --- | --- | --- | --- | --- |
| Height | 1410 ± 262a | 1180 ± 289a | 1162 ± 213a | 1758 ± 497bc |
| Diameter | 4440 ± 678ad | 4077 ± 918ad | 2779 ± 291a | 5158 ± 65bc |
| Parameters ($\mu\text{m}$) | CP2 | CN1 | CN2 | P |
| Height | 1933 ± 547bc | 1833 ± 436bc | 1910 ± 387bc | 0.0001 |
| Diameter | 5300 ± 717b | 4120 ± 699a | 4600 ± 617c | 0.0001 |

(P = <0.05 significance) different letters in the same line indicates significance.

Larvae exposed to BTI in CL_50_ 0.37ppm e CL_90_ 0.06ppm presented a tendency of decreasing the height of the tubule in CP1 and increasing in CP2, but it was not significant for CN1 and CN2. However, the diameter presented a significant increase (p<0.0001) in CP1 and CP2 in comparison with the other experimental groups. Degenerate or necrosis cells were observed with cell content leaking into the tubular lumen.

The Malpighi tubules of larvae in CN1 and CN2 groups did not present injuries in the abdominal region, besides presenting large and elongated nuclei, abundant basophilic cytoplasm with numerous mineralized cytoplasmic concretions of varying sizes for the storage of ions, metals and uric acid, and a dense matrix of elongated villi forming the brush border in the tubular lumen (Figure 3).

**Figure 3:**
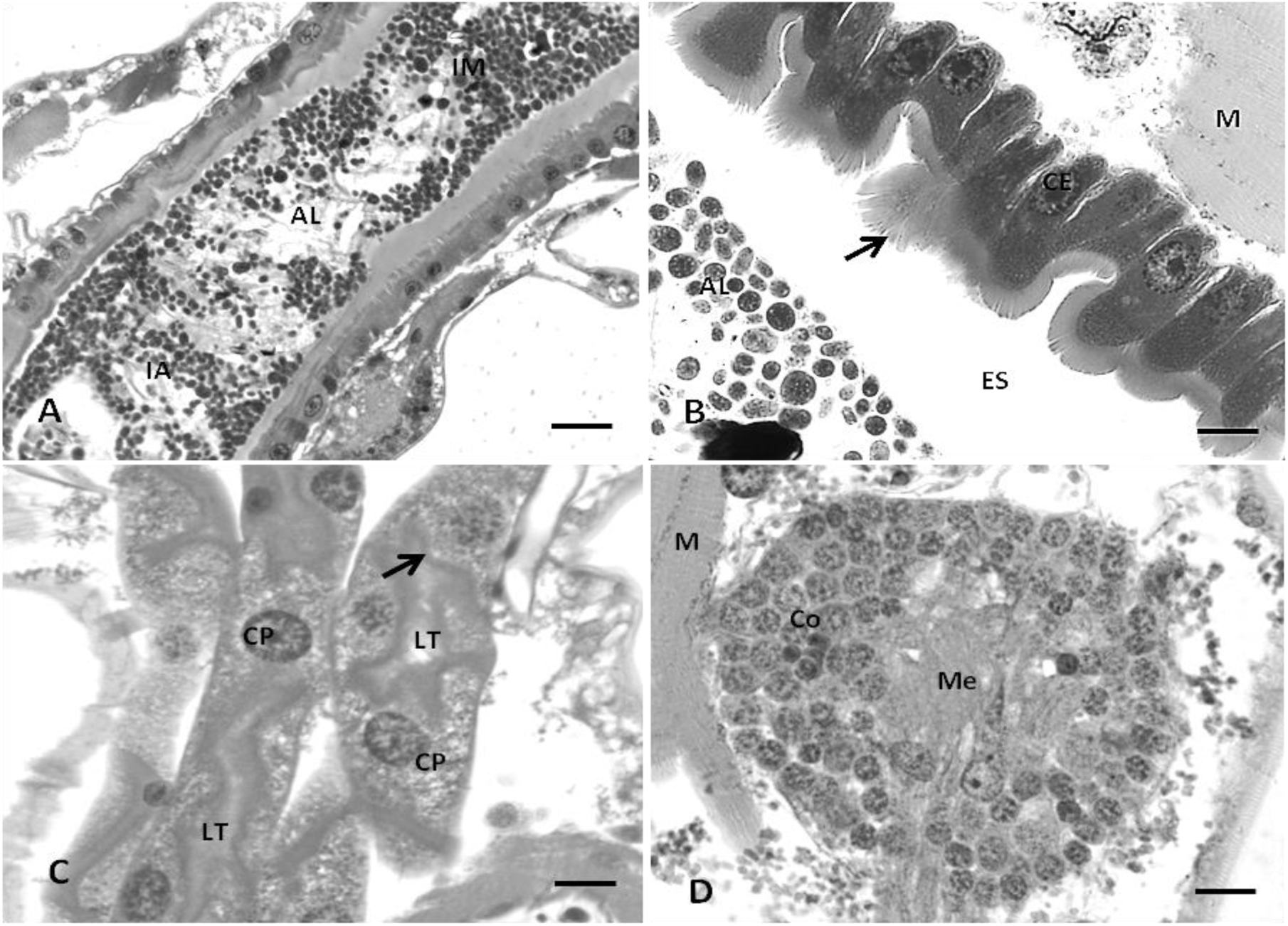
Photomicrography of intestine, Malpighi tubule and ganglia of third stage larvae of *Aedes aegypti* from the negative control group during a 24-hour observation. A and B: Intestine of CN1 with preserved structures. 50µm bar. B: Malpighi tubules of CN1. 50µm bar. C: Ganglia of CN1. 50µm bar. Anterior intestine (IA); Midgut (IM); Food (AL); Brush border (arrow); Sub peritrophic space (ES); Muscle (M); Main cell (CP); tubular lumen (LT); Ganglia’s cortical region (Co); Ganglia’s medullar region (Me).

### Ganglia

The nervous ganglia of G1, G2, and G3 are shown in Figure 1. It is observed that regardless of the product concentration, there was cortical necrosis and vacuolization of the basal ganglia, which were most observed in G3. Also, there was degeneration and diffuse necrosis with loss of structural organization in the positive control groups (Figure 2), while in the negative control groups there was no structural changes (Figure 3). These presented a cortical region formed by neuron bodies and a medullary region with a prothoracic gland and amorphous mass.

## Discussion

The control of mosquitoes is needed to improve health and quality of life in development countries due to the absence of general knowledge, development of resistance and socioeconomic reasons (Muthukumaran et al., 2015). The BTI toxicity is drastically reduced for mosquitoes due to a strong decrease in Cyt bioavailability. The low environmental persistence of Cyt in comparison with Cry toxins can lead to changes in the proportion of Cyt/Cry toxins persisting in the mosquitoes’ creation sites, besides facilitates the development of resistance to Cry toxins in exposed mosquitoes (Stalinski et al., 2014). Thus, new alternatives against the development of *Aedes aegypti* larvae is needed.

The *A. indica* oil present a toxic effect against *Anopheles gambiae* larvae, in addition to inhibiting pupal development (Okumu et al., 2007). Nano capsules with *M. alternifolia* oil have larvicide potential against *A. aegypti* (PIRES, 2019). The Andirobin present in *Carapa guianensis* causes apoptosis of the intestines of *A. aegypti* and *A. albopictus* larvae, causing their death within hours after ingestion (SILVA et al., 2006). In addition, Rawani et al. (2009) tested the aqueous seed extract of *C. papaya* and verified its larvicide activity against *Culex quinquefasciatus*.

The product formulated based on *Azadirachta indica, Melaleuca alternifolia*, and *Carapa guianensis* oils and *Carica papaya* bacterial fermented extract presented a greater toxic effect on the midgut of *Aedes aegypti* larvae in lower concentrations, diverging from Simon (2016) and Arruda et al., (2003), which tested *Eucalyptus staigeriana* and *Magonia Pubescens*, respectively.

In isolation, some of the compounds of the formulated product evaluated here present larvicide effect, but the effectiveness of the combination of these compounds was not previously tested. Here the toxicity of the formulated product on for the mesentery of *A. aegypti* larvae was noticed, mainly through the extrusion of the peritrophic matrix, since it is a defense mechanism of intestinal cells to eliminate the toxic intestinal content (Arruda et al., 2003; Barreto et al., 2006; VALOTTO et al., 2010, Valotto et al., 2011; GUIMARÃES, 2014).

These observations corroborate several authors that tested the effect of *Persea americana, Alnus glutinosa, Populus nigra, Quercus robur, M. pubescens*, and *Copaifera* extract and *Carapa guianensis* oil on *Aedes aegypti* (Arruda et al. 2003a; Abed et al., 2007; Silva et al., 2004; Valotto et al. 2010; Guimarães 2014).

The distal segment of the malpighi tubule of *A. aegypti* is constituted by main cells and star cells with a 5:1 distribution (Yu and BEYENBACH, 2004; Beyenbach et al., 2009, Beyenbach et al., 2010). These transepithelial main cells occupy about 90% of the tubule (Wu and Beyenbach, 2003) and are responsible by mediating the secretion of sodium ions (Na^+^) and potassium (K^+^), through active transport from hemolymph to tubular lumen (Beyenbach, 2003; Yu and Beyenbach, 2004). Thus, it is possible that the ionic transport has been compromised due to the injuries found in these cells of the evaluated larvae.

According to literature, it is know that the *Copaifera* oil causes apoptosis with partial or total destruction of epithelial cells, cytoplasmatic vacuolization, increase of sub peritrophic space with internal accumulation of acidophilic material, and damage to microvilli and peripheral nerves of *A. aegypti* larvae (Abed et al., 2007).

Similar injuries to those previously described were verified in groups CP1 and CP2, although for these larvae the damage was greater in anterior and posterior intestine and in the gastric cecum. This can occur due to the toxin-binding sites present in these organs (LIMA et al., 2003; Ruiz et al., 2004). The *Bacillus thuringiensis* cause changes in the ionic balance of intestinal cells and basal labyrinth, resulting in the increase of the cell surface or intercellular space and excessive transport of cell fluid to hemocoel (Bauer and Pankratz, 1992).

Although there are few reports in the literature regarding structural changes in ganglia of *A. aegypti* promoted by synthetic or natural molecules, there are studies indicating that azadiractin and meliantrium, components of *A. indica*, can inhibit endocrine glands that controls the metamorphosis and larval development, as well as degenerate the *corpus cardiacum* of the larvae, leading to a decrease in neurosecretory material (Valladares et al., 1997). This degenerative process promotes a decrease in morphogenetic hormonal levels in larvae and young insects (Viegas Junior, 2003).

Thus, it was found that the exposure of the *A. aegypti* third stage larvae to the formulated product composed by *Azadirachta indica, Melaleuca alternifolia*, and *Carapa guianensis* oils and *Carica papaya* fermented extract, at concentrations of 25 and 12.5%, caused irreversible cell damage in the larvae’s mesentery, Malpighi tubule and ganglia. Thus, we conclude that the formulated product can be used in the populational control of *Aedes aegypti* larvae, although studies focusing on field populations and larvae in different stages are still needed.

